# LARP4 Is an RNA-Binding Protein That Binds Nuclear-Encoded Mitochondrial mRNAs To Promote Mitochondrial Function

**DOI:** 10.1101/2022.10.24.513614

**Authors:** Benjamin M Lewis, Chae Yun Cho, Hsuan-Lin Her, Tony Hunter, Gene W Yeo

## Abstract

Mitochondrial associated RNA-binding proteins (RBPs) have emerged as key contributors to mitochondrial biogenesis and homeostasis. With few examples described, we set out to identify RBPs that regulate nuclear-encoded mitochondrial mRNAs (NEMmRNAs). Our systematic analysis of RNA-targets of 150 RBPs identified RBPs with a preference for binding NEMmRNAs, including LARP4, a La RBP family member. We show that LARP4’s targets are particularly enriched in mRNAs that encode respiratory chain complex proteins (RCCPs) and mitochondrial ribosome proteins (MRPs) across multiple human cell lines. Quantitative proteomics of cells lacking LARP4 show that protein levels of RCCPs and MRPs are significantly reduced. Furthermore, we show that LARP4 depletion reduces mitochondrial function, and that this phenotype is rescued by LARP4 re-expression. Our findings shed light onto a novel function for LARP4 as an RBP that binds to NEMmRNAs to promote mitochondrial respiratory function.

**Highlights:**

- Analysis of CLIP data reveals RBPs with a preference for mitochondrial mRNA targets
- LARP4’s RNA-target set is enriched for OXPHOS and mitochondrial ribosomal proteins
- Loss of LARP4 reduces protein levels of these two groups of mitochondrial proteins
- LARP4 is required for normal proliferation, translation, and OXPHOS function

## Introduction

A hallmark of eukaryotic cells is the presence of membrane-bound organelles, which facilitate the compartmentalization of biological processes. The protein components of these organelles are frequently recruited to them by polypeptide sorting sequences contained within the proteins that require localization. These cis-acting polypeptide sequences are recognized by other effector proteins that facilitate subcellular localization by various mechanisms (Bolender et al., 2008). Interestingly, an additional pathway for protein targeting is mediated by information contained within the mRNA sequence that encodes the protein. These cis-acting mRNA sequences are recognized by RNA-binding proteins (RBPs), which in turn influence the subcellular distribution and translational dynamics of their bound mRNA transcripts to promote synthesis of proteins in the proximity of their functional locations (Bethune et al., 2019).

Efficient protein targeting is particularly important for mitochondrial proteins, many of which have biochemical or biological properties that are detrimental to the cell outside of the context of the mitochondrion (Yoo et al., 2005, Boos et al., 2019, Bykov et al., 2020, Williams et al., 2014, Wrobel et al., 2014). The classical pathway for protein targeting to the mitochondria is polypeptide mediated, facilitated by an N-terminal mitochondrial targeting sequence present on many but not all mitochondrial precursor proteins (Bolender et al.,2008). Additionally, several mRNA-mediated mitochondrial targeting pathways have been described in lower eukaryotes and higher eukaryotes (Bethune et al., 2019, Bykov et al., 2020). Many of the RBPs and interacting proteins involved in these mRNA-mediated pathways were first described in yeast or Drosophila (Eliyahu et al., 2010, Fields et al., 1998, Sen. et al., 2015, Zhang et al., 2016) and were later found to have a conserved or similar function in human cells (Gabrovsek et al., 2020, Gao et al. 2014, Gehrke et al., 2015, Matsumoto. et al., 2014, Schatton et al. 2017). Some factors involved in mitochondrial mRNA-dependent targeting described in yeast, however, do not have a human homolog with a conserved function (Garcia-Rodriguez et al., 2007, Saint-Georges et al., 2008, Lesnik et al., 2014, Zabezhinsky et al., 2016). The extent to which human RBPs regulate expression of nuclear-encoded mitochondrial mRNAs (NEMmRNAs) is not fully understood.

With the goal of identifying human RBPs with a novel role in mitochondrial biology, we made use of the ENCODE collection of 223 enhanced CLIP (eCLIP) datasets profiling 150 RBPs in K562 and HepG2 cell lines in a standardized workflow (Van Nostrand et al, 2020). Through a systematic computational analysis of these eCLIP datasets, several RBPs with RNA-target sets enriched for NEMmRNAs were identified. Of these NEMmRNA-enriched RBPs, LARP4, a La-related RBP, was selected for further study, due to an enrichment for RNA-targets encoding respiratory chain complex proteins (RCCPs) and mitochondrial ribosome proteins (MRPs). LARP4 was first identified as one of the La-related proteins (LARPs), each of which are paralogs to the conserved La protein, which functions as a processing chaperone for RNA transcripts produced by RNA polymerase III (Bousquet-antoneli et al., 2009). The La protein and the LARPs each contain an evolutionarily related RNA-binding protein domain called the La module. However, the La modules found on LARPs have diverged to bind RNA sequences distinct from their ancestral paralog. Additionally, many of the LARPs have gained additional protein domains to facilitate novel protein-protein interactions (Maraia et al., 2017). Upon initial characterization, LARP4 was shown to bind directly to poly-A RNA as well as indirectly through protein-protein interactions with poly-A binding proteins (PABPs) (Yang et al., 2011). Additionally, LARP4 was shown to interact with the 40S ribosomal protein RACK1 and assemble into translating polysomes (Yang et al., 2011). Later studies demonstrated that LARP4 plays a functional role in maintaining poly-A tail length, likely through competition with deadenylases for PABP binding sites (Mattijssen et al., 2017, and Mattijssen et al., 2020). Recently, LARP4 was reported to be recruited to the mitochondrial surface through an interaction with the PKA adaptor protein AKAP1, which also functions as an RNA-binding protein (Gabrovsek et al., 2020). Without knowledge of the specific RNA targets of LARP4, those authors proposed LARP4 acted as an RNP scaffolding factor which could facilitate the recruitment of RNP-associated translation factors to mitochondria to enhance local translation of mRNAs bound by AKAP1 at the mitochondrial surface (Gabrovsek et al., 2020). In this study, we systematically determine the RNA targets of LARP4 to expand this model to support a role for LARP4 as transcript-specific NEMmRNA recruitment factor.

Here, we identified and mapped the binding sites that LARP4 has on 713 human mRNA targets, of which 186 or 26% are transcripts that encode mitochondrial proteins, with enrichment in proteins involved in oxidative phosphorylation (e.g., RCCPs) or mitochondrial translation (e.g., MRPs). Additionally, we showed that CRISPR-mediated LARP4 depletion results in reduced abundance of the steady state levels of these two groups of proteins. Furthermore, LARP4-depleted cells show reduced oxidative phosphorylation capacity, and this phenotype is ameliorated by re-expression of LARP4. Our results indicate that LARP4 is an RBP that binds NEMmRNAs and promotes the expression of proteins essential to mitochondrial respiratory function to enhance respiratory function.

## Results

### LARP4 has a preference for binding nuclear-encoded mitochondrial mRNAs

The ENCODE consortium has recently made publicly available the RNA-binding profiles of hundreds of RNA binding proteins (RBPs). This was done by performing the eCLIP-seq assay for 150 RBPs in two human cell lines (HepG2 hepatocellular carcinoma and K562 erythroleukemia) for each RBP in duplicates (Van Nostrand et al, 2020). We analyzed these ENCODE eCLIP datasets for RBPs whose RNA-target sets were enriched for mRNAs encoding either proteins localized to the mitochondria (Mitocarta_2.0) or proteins specifically involved in oxidative phosphorylation (KEGG OXPHOS pathway) (Figures 1A, 1C and 1D), and identified several RBPs with RNA-target sets substantially enriched for both: LARP4, DDX3X, RPS3, SUB1, PABPC4, YBX3 and several others (Figures 1C and 1D). Among them, LARP4 showed the greatest enrichment, and was chosen for more in-depth analysis. We performed an independent LARP4 eCLIP sequencing experiment in human HEK293 embryonic kidney cells (Figure 1B) and discovered an even greater enrichment for RNA-targets encoding mitochondrial or oxidative phosphorylation proteins than observed in any of the other ENCODE eCLIP datasets available at the time (Figures 1C and 1D).

**Figure 1.**
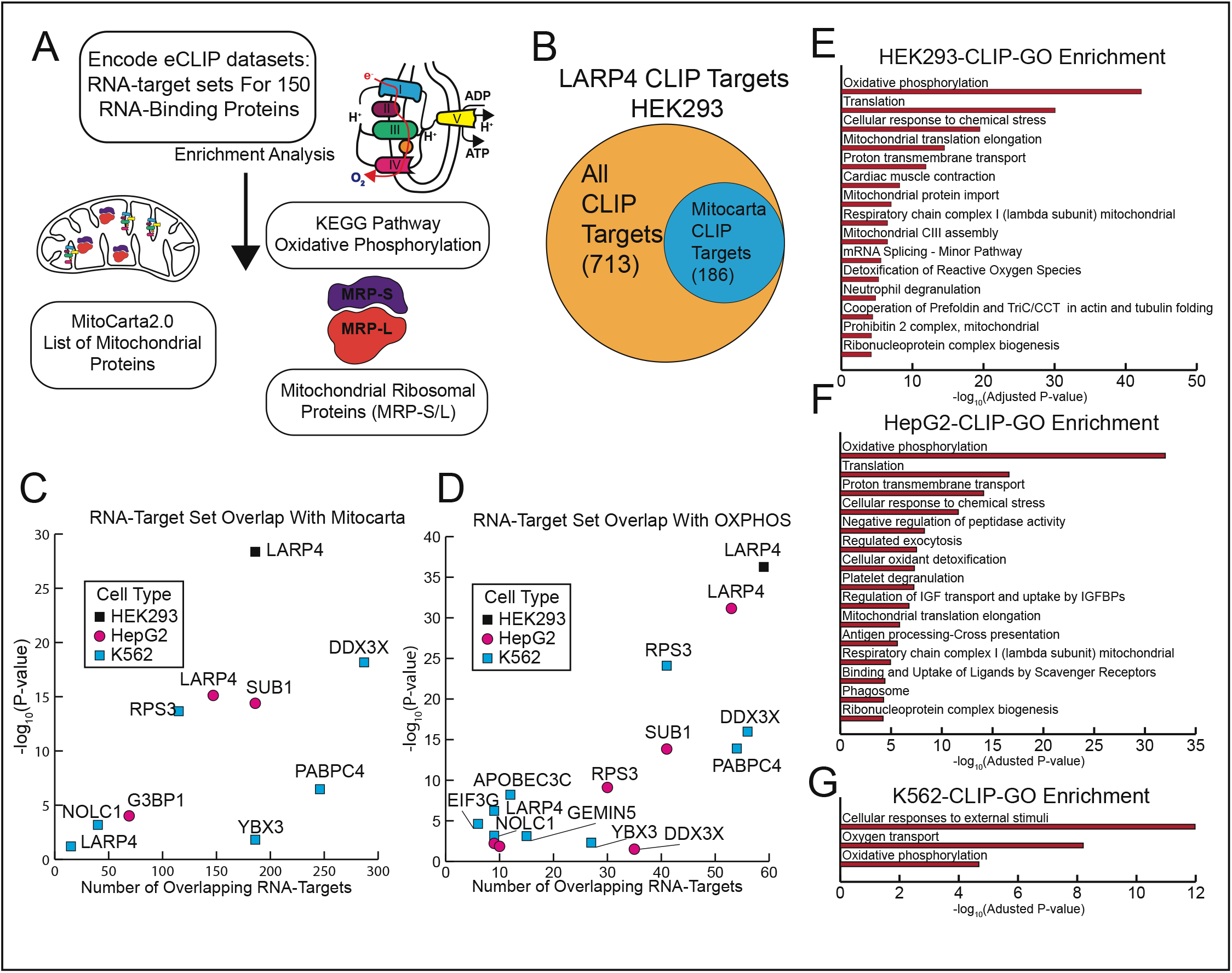
LARP4 has a preference for binding nuclear-encoded mitochondrial mRNAs. (A) Overview of enrichment analysis performed on eCLIP dataset for 150 RBPs in K562 and HepG2 cell lines. (B) Diagram showing the proportion of LARP4 CLIP targets that encode for mitochondrial proteins (C) Enrichment analysis results for RNA-target set (eCLIP data) overlap with mitochondrial genes. The hypergeometric test for significance of overlap for each RBP dataset is plotted on the y-axis and the number of overlapping targets is plotted on the x-axis. RBPs with significant overlap (p-value≤0.05) with LARP4-K562 also shown (p-value=O.06). (D) Enrichment analysis results for RNA-target set (eCLIP data) overlap with proteins in the KEGG oxidative phosphorylation pathway (HSA_00190). Only RBPs with significant overlap (p-value≤0.05) are shown. (E-G) Metascape gene ontology analysis performed on the three LARP4 CLIP-seq data sets from HEK293 (E), HepG2 (F), K562 (G) cells. Gene-target lists of each CLIP-seq data set are defined as genes encoding an mRNA containing at least one CLIP peak that passes significance thresholds (-log10Pvalue > 7, log2foldchange (IP/input) > 4). See also Figure S1.

Overlap between the gene-target lists, defined as genes encoding an mRNA containing at least one significant LARP4 CLIP peak, was greatest between the LARP4 eCLIP dataset generated from HEK293 and HepG2 cells, which are both of epithelial origin, with the K562 cell dataset generating fewer overall gene-targets (Figure S1C). Gene ontology analysis was performed on these eCLIP datasets, using similarly defined gene-target lists as a foreground and dataset specific background gene lists using the Metascape method (Figures 1E, 1F and 1G) (Zhou et al., 2019). Oxidative phosphorylation (GO: 0006119) was the most enriched gene ontology term for both the HEK293 and HepG2 cell datasets (Figures 1E and 1F) and one of the top three terms for the analysis performed on the K562 eCLIP dataset (Figure 1G). The overlap between gene targets of each LARP4 eCLIP dataset was even greater when only considering CLIP targets that are also mitochondrial mRNAs (Figure S1D).

The translation (RHSA:72766) gene ontology term, which includes proteins that make up the cytosolic ribosome as well as the mitochondrial ribosome, was the second most enriched term in both the HEK293 and HepG2 cell eCLIP datasets. This enrichment for translational genes was also substantial when compared to other RBPs in the ENCODE dataset collection (Figure S1B). Furthermore, mitochondrial translation elongation (RHSA-5389840), a term that primarily composed of genes that encode mitochondrial ribosome proteins (MRPs), was also a top gene ontology term of both the HEK293 and HepG2 cell eCLIP datasets. (Figures 1E and 1F). A separate analysis looking for overlap between the genes that encode LARP4’s mRNA-targets and the genes that encode the MRPs (GO:OOO5761) confirms that MRPs are highly enriched (p-value=6e-11) in the HEK293 LARP4 eCLIP target set (Figure S1F). This enrichment for MRPs was also substantial when compared to other RBPs in the ENCODE dataset collection (Figure S1A). A similar overlap analysis showed that genes encoding proteins that function in the oxidative phosphorylation pathway i.e., respiratory chain complex proteins (RCCPs) are also highly enriched (p-value=5e^-37^) in the LARP4 eCLIP target set (Figure S1E).

How LARP4 achieves its RNA-target specificity is not fully understood. Using the eCLIP datasets, we performed k-mer analysis to identify possible linear sequence motifs in LARP4’s RNA-target sets. While robust motifs were not identified, there was a weak degenerate C-rich motif present within the LARP4 peaks present in 3’ UTRs of target genes (data not shown). We also performed a similar k-mer analysis on the following subsets of LARP4 targets, nuclear-encoded mitochondrial genes, OXPHOS genes and mitochondrial ribosome proteins genes. These searches also did not identify any robust motifs. Metagene analysis of LARP4’s eCLIP data shows that LARP4 coats many of its target mRNAs from CDS to 3’ UTR with the highest peak density centered around the stop codon and a small peak of density around the start codon; this pattern was consistent across subsets of LARP4 targets (Figures S1G), perhaps explaining why there is an absence of specific strong motifs. LARP4 contains both RNA-binding domains (La-module) and protein interaction domains that interact with other RBPs or RNA-associated proteins such as Poly-A binding proteins (PABPs) and RACK1 (Maraia et al., 2017). It is possible that LARP4’s RNA-target specificity is achieved through a cooperative mechanism involving both its RNA-binding domains and its protein interaction domains.

Together these data identify LARP4 as an RNA-binding protein with a specific preference for nuclear-encoded mitochondrial mRNAs (NEmRNAs). Of these NEmRNAs targets, LARP4 has a particular preference for binding mRNAs encoding RCCPs as well as MRPs (Figures 1E and 1F).

### Loss of LARP4 disrupts protein levels without affecting mRNA abundance of mitochondrial targets

To study the functional consequences of LARP4 depletion, knockout cell (KO) lines were generated for HEK293 cells and the U2OS human osteosarcoma cell line. Genome editing was performed by transient expression of CRISPR/Cas9 and guides targeting the first exon of LARP4 and the upstream promotor region. Disruption of LARP4 expression was validated by immunoblot analysis (Figures 2B and 2C), and sequencing of genomic regions surrounding the CRISPR guide sites from the KO cell line subclones confirmed disruption of all LARP4 alleles (Figure S2).

**Figure 2.**
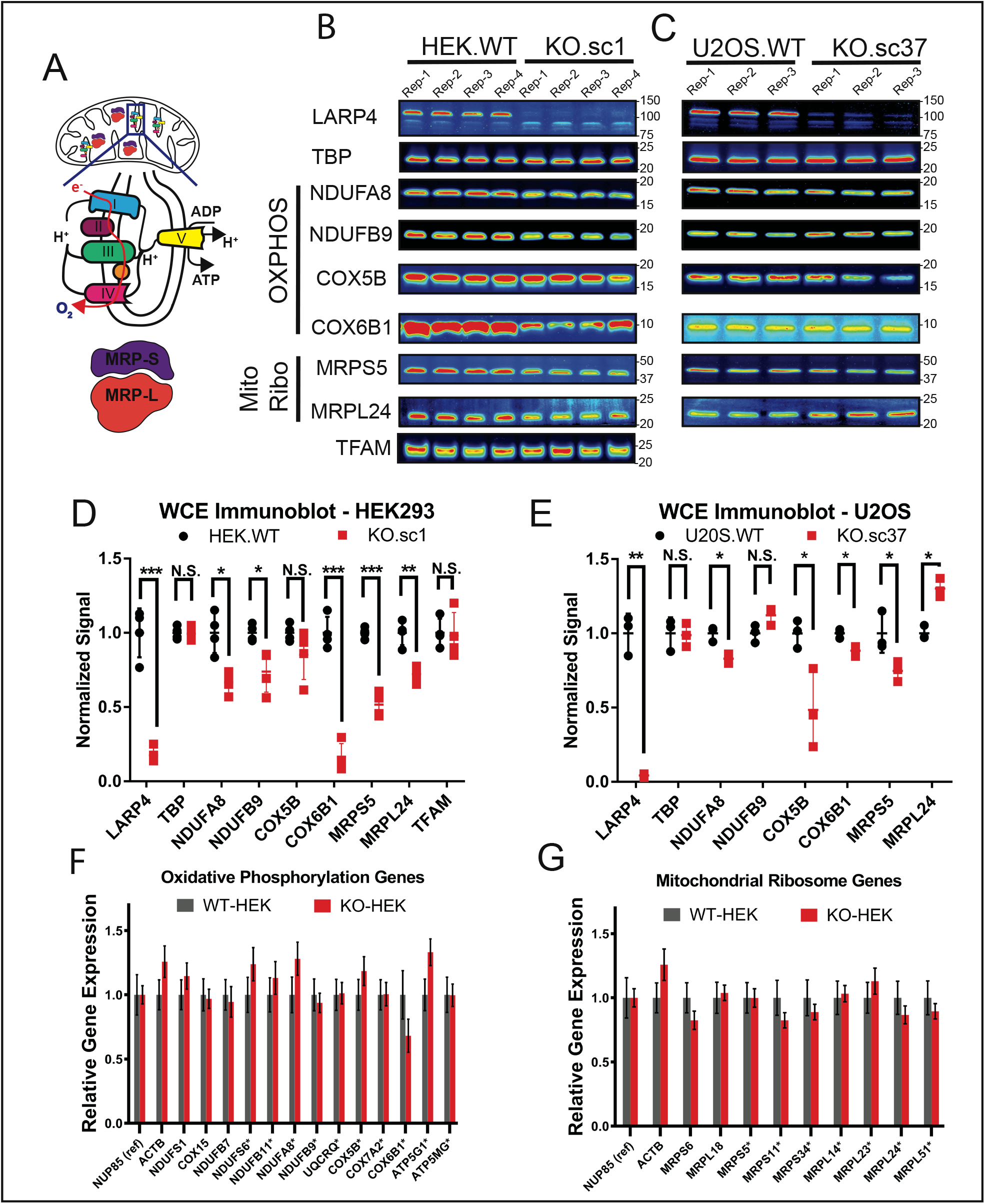
Loss of LARP4 disrupts protein levels without affecting mRNA abundance of mitochondrial targets. (A) Diagram of the mitochondrial, oxidative phosphorylation proteins and mitochondrial ribosome proteins. (B-C) Validation of LARP4 depletion by CRISPR/Cas9 by immunoblot in the clonal HEK293 cell line (B) and the clonal U2OS cell line (C). Immunoblot analysis of various oxidative phosphorylation proteins and mitochondrial ribosome proteins in the HEK293 KO and parental WT cells (B) and in the U2OS KO and parental WT cells (C). (D-E) Quantification of biological replicates shown in immunoblot panels. Band intensities are normalized by a quantitative total protein stain. (F-G) Analysis of mRNA abundance in the HEK293 KO/WT cells by qPCR using probes targeting mitochondrial ribosome subunits proteins (F) or oxidative phosphorylation proteins (G). LARP4 targets are denoted with an asterisk. All averages shown are from independent biological replicates. See also Figure S2.

Because mRNAs encoding RCCPs and MRPs were particularly enriched in LARP4’s RNA target set, expression levels of representative proteins from these protein complexes were characterized by immunoblot analysis (Figures 2A-2E). Based on the strong LARP4 binding to their encoding transcripts NDUFA8, NDUFB9, COX5B and COX6B1 were selected as representative RCCPs; MRPS5 and MRPL24 were selected as representative MRPs. The TATA-Box binding protein (TBP) was included as a non-LARP4 target loading control. Biological replicate samples of whole cell extracts were collected from KO cell lines and their respective wildtype parental lines for analysis. Antibody signals were normalized with a quantitative total protein stain and replicate averages quantified (Figures 2D and 2E). Samples from HEK293^LARP4-/-^ KO cells (Figures 2B and 2D) showed a significant reduction in protein expression levels of multiple RCCPs (NDUFA8: KO/WT=0.67, NDUFB9: KO/WT=0.74 and COX6B1: KO/WT=0.16) and multiple MRPs (MRPS5: KO/WT=0.52 and MRPL24: KO/WT=0.72). Parallel analysis of samples from U2OS^LARP4-/-^ KO cells (Figures 2C and 2E) showed that protein levels were reduced in most cases (NDUFA8: KO/WT=0.83, COX5B: KO/WT=0.48, COX6B1: KO/WT=0.88 and MRPS5: KO/WT=0.75), but significantly increased in one case (MRPL24: KO/WT=1.30). The overall level of target protein disruption observed by immunoblot was less pronounced in the U2OS LARP4 knockout line. We also depleted LARP4 in HEK293 cells by transduction with a lentivirus vector expressing an shRNA targeting LARP4 (LARP4: shKD/NT-Con=0.71). Depletion of LARP4 by this orthogonal method also resulted in significant reduction in protein expression levels of OXPHOS proteins (NDUFA8: KO/WT=0.71, NDUFB9: KO/WT=0.85) and a representative MRP (MRPS5: KO/WT=0.76) (Figures S2D and S2E). These data show that loss of LARP4 results in disruption of protein levels of select proteins belonging to the RCCP and MRP gene groups that were enriched in LARP4’s RNA target set in two different human cell lines.

Given that LARP4 is thought to stabilize the mRNA levels of a subset of targets in certain contexts (Mattijssen et al. 2020, Yang et al., 2011), we sought to determine if the reduction in protein levels of RCCPs and MRPs observed in the HEK293^LARP4-/-^ KO cells could be explained by a corresponding reduction in abundance of their encoding mRNAs. Analysis of mRNA abundance was performed by qPCR on a panel of mRNAs encoding MRPs (Figure 2F) as well as RCCPs (Figure 2G) on wild-type and HEK293^LARP4-/-^ KO cells. None of the 9 mRNAs encoding MRPs tested or the 13 mRNAs encoding RCCPs tested showed a significant change in abundance relative to wild-type HEK293 cells, indicating that the reduction in protein levels of RCCPs and MRPs observed in the HEK293^LARP4-/-^ KO cells is not due to reduction in the abundance of the encoding mRNAs. These data show that the loss of LARP4 does not affect the levels of these target transcripts, suggesting that LARP4 is regulating the translation of these targets possibly by promoting localized translation in the vicinity of the mitochondria.

To assess the effect of LARP4 depletion on mitochondrial DNA expression we measured relative mRNA abundance of a panel of genes encoded on the mitochondrial chromosome. We did not find any significant changes in the HEK293^LARP4-/-^ KO cells relative to the wild-type HEK293 cells (Figure S2A). We also measured relative mRNA abundance of a panel genes regulated by the PGC-1α/NRF axis that controls TFAM expression and mitochondrial biogenesis. For the majority of these genes, we did not find any significant changes (Figure S2B). We also did not observe a significant change in TFAM protein expression by immunoblot (Figure 2B). Together these data show that loss of LARP4 does not result in changes in mitochondrial DNA levels or expression of TFAM.

### Quantitative proteomic analysis of LARP4 KO cell line reveals reduced abundance of mitochondrial ribosome proteins and OXPHOS proteins

To gain more comprehensive insights into the proteomic consequences of LARP4 depletion and verify the reductions in protein abundance observed for RCCPs and MRPs by immunoblotting using an orthogonal method of measurement, quantitative tandem mass tagging (TMT) proteomics was performed on HEK293^LARP4-/-^ KO and wild-type cells. To gain insights into possible defects in protein localization to the mitochondria in the HEK293^LARP4-/-^ KO cells subcellular fractionation was performed in parallel. A rapid-magnetic mitochondrial enrichment strategy was used to generate two types of protein extracts for each genotype, a whole-cell extract (WCE) and mitochondrial fraction extract (MITO) from the same cultures. Biological replicates (N=4) of each type of protein extract were processed and analyzed by separate quantitative TMT proteomic experiments, to produce a WCE-TMT dataset (Figure 3A) and a MITO-TMT dataset (Figure 3B).

**Figure 3.**
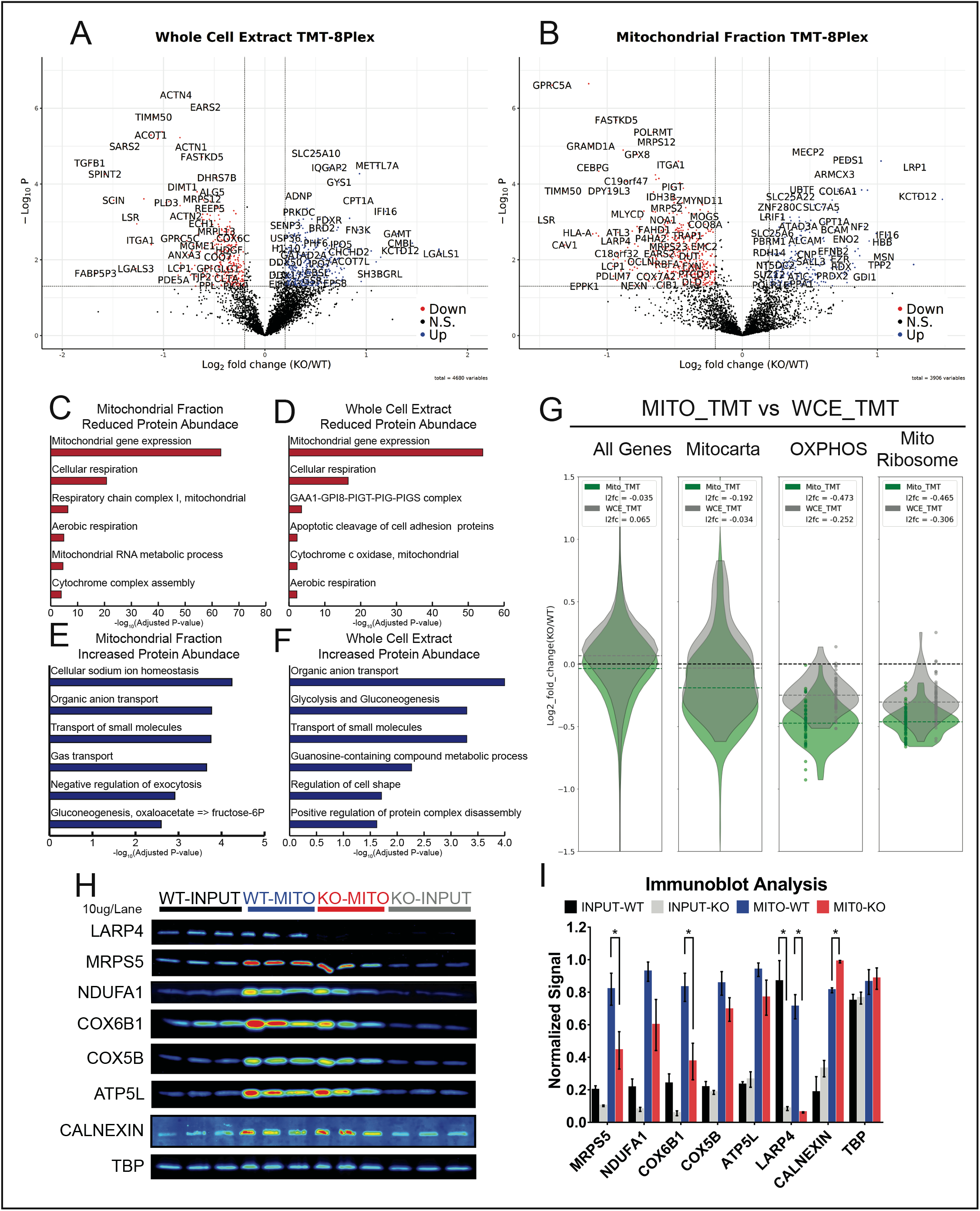
Quantitative proteomic analysis of LARP4 KO cell line reveals reduced abundance of mitochondrial ribosome proteins and OXPHOS proteins. (A) Volcano plot showing proteins abundance (tag ratio of KO/WT) results from quantitative tandem mass tagging (TMT) analysis of whole cell extracts (N=4) prepared form HEK293^LARP4-/-^ KO cells and wild-type cells. (B) Volcano plot showing proteins abundance results from quantitative tandem mass tagging (TMT) analysis of mitochondrial enriched extracts (N=4). (C-D) Metascape gene ontology analysis of proteins with significant decreases in abundance identified by TMT analysis in whole cell extracts (C) and mitochondrial enriched extracts (D). (E-F) Metascape gene ontology analysis of proteins with significant increase in abundance identified by TMT analysis in whole cell extracts (E) and mitochondrial enriched extracts (F). (G) Violin plots comparing the changes in protein abundance of OXPHOS proteins and mitochondrial ribosome proteins observed in the mitochondrial extract (green) and whole cell extract (grey) proteomics experiments. Distributions shown are of average log2 fold changes (KO/WT) of proteins within each gene group. (H) Immunoblot analysis of whole cell extracts (input) and mitochondrial enriched extracts, each lane was load with 10 ug of protein. (I) Quantification of immunoblot analysis. Significance thresholds for changes in protein abundance in TMT data are defined as P-value<0.05 and log2foldchange (KO/WT) +/- 0.2.

To determine how gene groups that were overrepresented within LARP4’s RNA-target set were affected by LARP4 depletion gene ontology (GO) analysis was performed using the Metascape method (Zhou et al., 2019). For each TMT dataset (WCE or MTIO), sets of proteins present in significantly increased (WCE: n=203, MITO: n=289) or decreased (WCE: n=286, MITO: n=494) abundance were defined (log2foldchange +/- 0.2 and p-value<0.05) and a separate GO analysis performed on each. The GO analysis of proteins with decreased abundance in HEK293^LARP4-/-^ KO cells (Figures 3C and 3D) revealed that many of the depleted proteins belonged to similar gene groups that were also overrepresented within LARP4’s RNA-target set. In both proteomic datasets, two of the most enriched gene groups in the set of depleted proteins were proteins involved in mitochondrial gene expression and proteins involved in cellular respiration. The mitochondrial gene expression term had 104 proteins present in the set of depleted proteins, primarily MRPs, and the cellular respiration term had 60 proteins present almost entirely RCCPs (Figures 3C and 3D). These gene groups were also enriched within LARP4’s RNA-target set (Figures 1E, S1E and S1F) suggesting that LARP4 binding enhances protein expression of these target mRNAs.

The one mitochondrially encoded protein detected in the TMT-proteomics dataset, MT-CO2, was depleted in the HEK293^LARP4-/-^ KO cells (WCE: KO/WT=0.86 and MITO:KO/WT=0.76) to levels similar to many of the nuclear encoded RCCPs, suggesting that either the depletion of MRPs results in decreased mitochondrial translation or other biological processes such as degradation or transcriptional buffering are occurring to bring the subunits of the respiratory complexes into stoichiometric balance. These same biological processes that keep the proportions of protein complex components in stoichiometric balance (Taggart et al., 2020) likely explain why the few non-LARP4 target RCCPs or MRPs present in the TMT-proteomics dataset are depleted to similar levels as the rest of the members of their protein complexes.

Fewer proteins with significantly increased abundance in HEK293^LARP4-/-^ cells were identified in both the WCE-TMT dataset (n=203) and MITO-TMT dataset (n=289). The GO analysis of these proteins with increased abundance in HEK293^LARP4-/-^ cells identified gene sets involved in biological processes that could conceivably compensate for reduced cellular respiration e.g., organic anion transport (WCE-TMT and MITO-TMT), glycolysis and gluconeogenesis (WCE-TMT), and transport of small molecules (WCE-TMT and MITO-TMT) (Figures 3E and 3F). Furthermore, these biological processes were not enriched in the LARP4 RNA-target eCLIP dataset (Figure 1E). The increase in abundance of proteins with potential to compensate for reduced cellular respiration, indicate the possibility that the HEK293^LARP4-/-^ KO cells have undergone some type of genetic or metabolic compensation for the loss of LARP4 and associated reduction in abundance of proteins involved in cellular respiration. To validate some of the changes in protein abundance observed in the TMT experiments, quantification of protein abundance by immunoblot analysis of selected RCCPs and MRPs was also performed on similar WCE extracts and MITO extracts produced while optimizing the mitochondrial enrichment strategy. Although increased variability in these pilot experiments reduced the significance of some comparisons, significant differences were observed (MRPS5 and COX6B1) as well similar downward trends (Figures 3H and 3I).

Interestingly when comparing the two TMT experiments, the reduction in protein abundance of RCCPs and MRPs was much more pronounced in the analysis of the mitochondrial extracts compared to the analysis of the whole cell extracts from the same cultures. For the OXPHOS proteins (RCCPs) the average fold change in protein abundance was 0.72 in the mitochondrial extracts and 0.84 in the whole cell extracts (Figure 3G). A similar trend was observed for the MRPs with an average fold change of 0.73 in the mitochondrial extracts and an average fold change of 0.81 in the whole cell extracts (Figure 3G). Suggesting that in the HEK293^LARP4-/-^ KO cells targeting of these proteins to the mitochondria is impaired. These observations support a model in which LARP4 promotes both the translation and subcellular targeting of RCCPs and MRPs to the mitochondria.

### Loss of LARP4 reduces cell proliferation rates, levels of oxidized proteins and translation rates

Given the essential role of mitochondria in cell proliferation (Birsoy et al., 2015 and Sullivan at al., 2015) and the depletion of proteins essential to mitochondrial function in observed in LARP4 KO cells, we hypothesized that cell proliferation would be reduced in LARP4 KO cells. Cell proliferation studies showed that the HEK293^LARP4-/-^ KO cells had significantly reduced proliferation rates, averages from independent experiments (N=3) showed significantly increased cell doubling times relative to wild type cells, and this reduction in the rate of cell proliferation was completely rescued by re-expression of LARP4 (Figure 4A). A reduced rate of cell proliferation was also observed in the U2OS^LARP4-/-^ KO cells compared to wild type cells in two independent experiments (Figure 4B). These data indicate that LARP4 is required for normal rates of proliferation in HEK293 cells and that LARP4 depletion is also associated with reduced proliferation rates in the cancer cell line U2OS.

**Figure 4.**
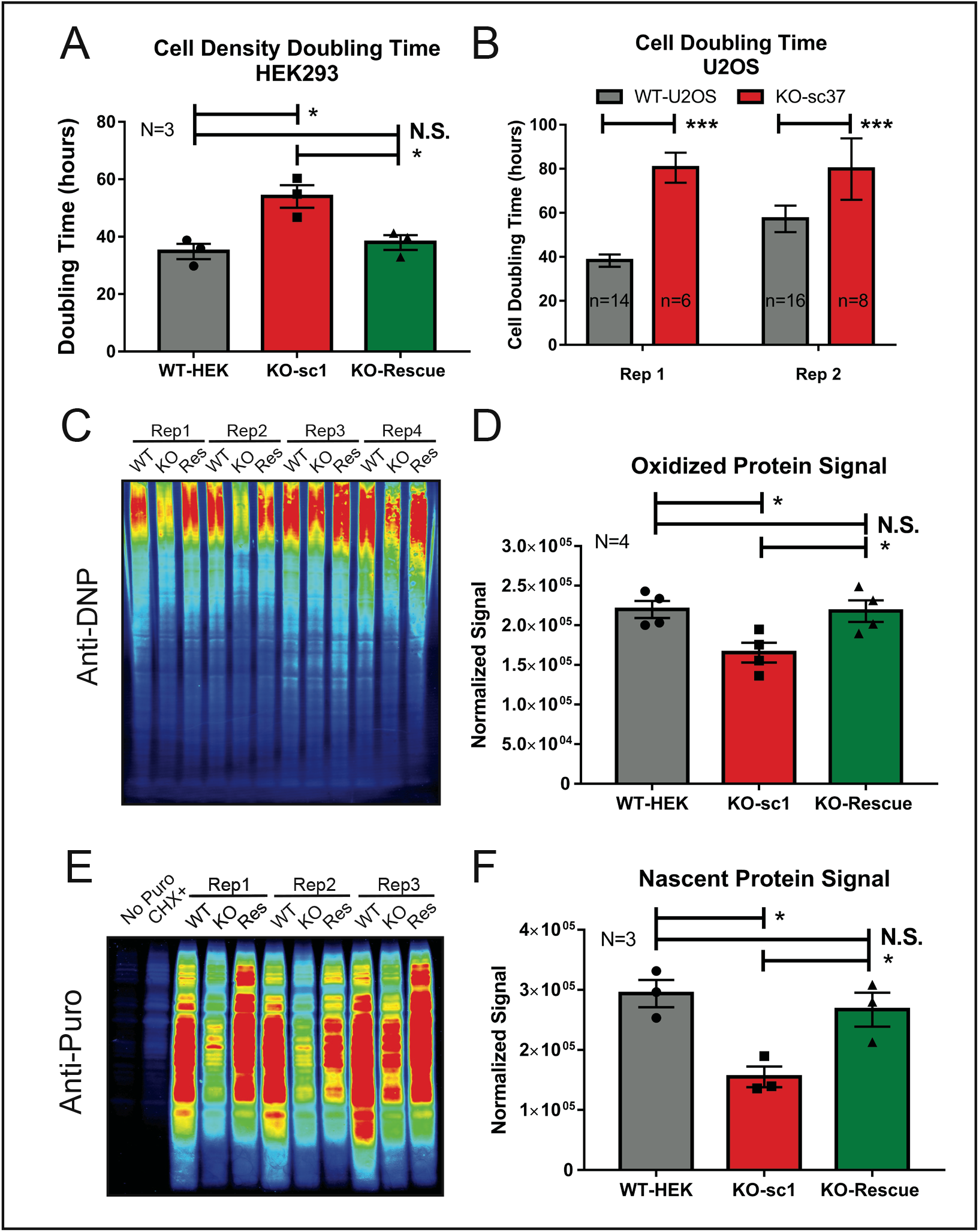
Loss of LARP4 reduces cell proliferation rates, levels of oxidized proteins and protein translation rates. (A) A summary plot of average confluency doubling times for wild-type HEK293 cells, HEK293^LARP4-/-^ KO cells and HEK293^LARP4-/-^ KO-Rescue cells from independent experiments (N=3). (B) A plot of cell doubling times of wild-type U2OS cells and U2OS^LARP4-/-^ KO cells, averages of technical replicates from each of two independent experiments are shown (Rep 1 and Rep 2). (C-D) Analysis of oxidized protein abundance by the oxiblot assay (C) and quantification of normalized oxidized protein signal averages (N=4) for each cell line, each replicate represents an independent experiment (D). (E-F) Analysis of translation rates by the puromycin incorporation assay (E) and quantification of normalized puromycin incorporation signal averages (N=3) for each cell line, each replicate represents an independent experiment (F). See also Figure S3.

Because mitochondria are a major source of reactive oxygen species (Balaban et al., 2005), which are a cause of protein oxidation, levels of oxidized proteins were measured, using a derivatization and immunoblot approach (Oxiblot Kit). This analysis found that oxidized proteins were present in significantly lower abundance in HEK293^LARP4-/-^ KO cells compared to wild type cells, and that this phenotype was completely rescued by re-expression of LARP4 (Figures 4C, 4D and S3A-S3D). These data indicate that LARP4 promotes the production of oxidized proteins, likely through promoting mitochondrial respiration rates.

LARP4 has been shown to associate with translational machinery (Yang et al., 2011) and to enrich mRNAs encoding translational proteins in its RNA-target set (Figures 1E, 1F and S1B). Additionally, protein translation is an energy intensive process, and LARP4 targets and regulates expression of proteins essential to the energy producing capacity of the mitochondria. For these reasons we wanted to assay translation rates, and for this purpose used the puromycin incorporation assay. The translation rates in the HEK293^LARP4-/-^ KO cells were found to be substantially reduced relative to the wild type cells as measured by the normalized puromycin incorporation signal. Additionally, re-expression of LARP4 returned translation rates to levels similar to wild type cells (Figures 4E, 4F and S3E-S3H). These data indicate that LARP4 is required for normal translation rates in HEK293 cells.

### LARP4 Promotes Oxidative Phosphorylation Function

As proteins essential for cellular respiration (RCCPs and MRPs) were significantly depleted in the HEK293^LARP4-/-^ KO cells, the functional impact on oxidative phosphorylation (OXPHOS) function and capacity was investigated. The Seahorse Extracellular Flux Analyzer was used to measure oxygen consumption rates (a parameter directly related to OXPHOS function). The analyzer was configured to run the Mito-stress-test assay which assesses basal respiration and maximal respiration, measured in the terms of oxygen consumption rates (OCR).

In every Mito-stress-test Seahorse assay performed, the HEK293^LARP4-/-^ KO cells exhibited significantly reduced basal respiration and maximal respiration compared to wild type cells (Figures 5A, 5B and S4A-S4C). Similarly, the U2OS^LARP4-/-^ KO cells exhibited deficiencies in OXPHOS function as measured by the Mito-stress-test Seahorse assay (Figures 5C, 5D and S5A-S5C). Furthermore, we demonstrate that rescue of LARP4 depletion by overexpression of LARP4 in HEK293^LARP4-/-^ KO cells significantly improved the OXPHOS deficiency in the LARP4-deficient cells although not completely to wild type levels (Figures 5A, 5B and S4A-S4C). It is likely that the LARP4 KO cells undergo compensatory metabolic changes following genetic disruption of LARP4 and these changes prevent a complete restoration of OXPHOS rates upon LARP4 reexpression. Evidence of compensatory metabolic change were observed in the analysis of the HEK293^LARP4-/-^ KO cells by TMT-proteomics e.g., upregulation of proteins involved in glycolysis and gluconeogenesis (Figures 3E and 3F). These results from two different human cell lines demonstrate that loss of LARP4 reduces oxidative phosphorylation rates, indicating that depletion of RCCPs and MRPs observed in LARP4 KO cells by TMT-proteomics and immunoblot analysis has a functional impact on the biological processes carried out by these protein complexes.

**Figure 5.**
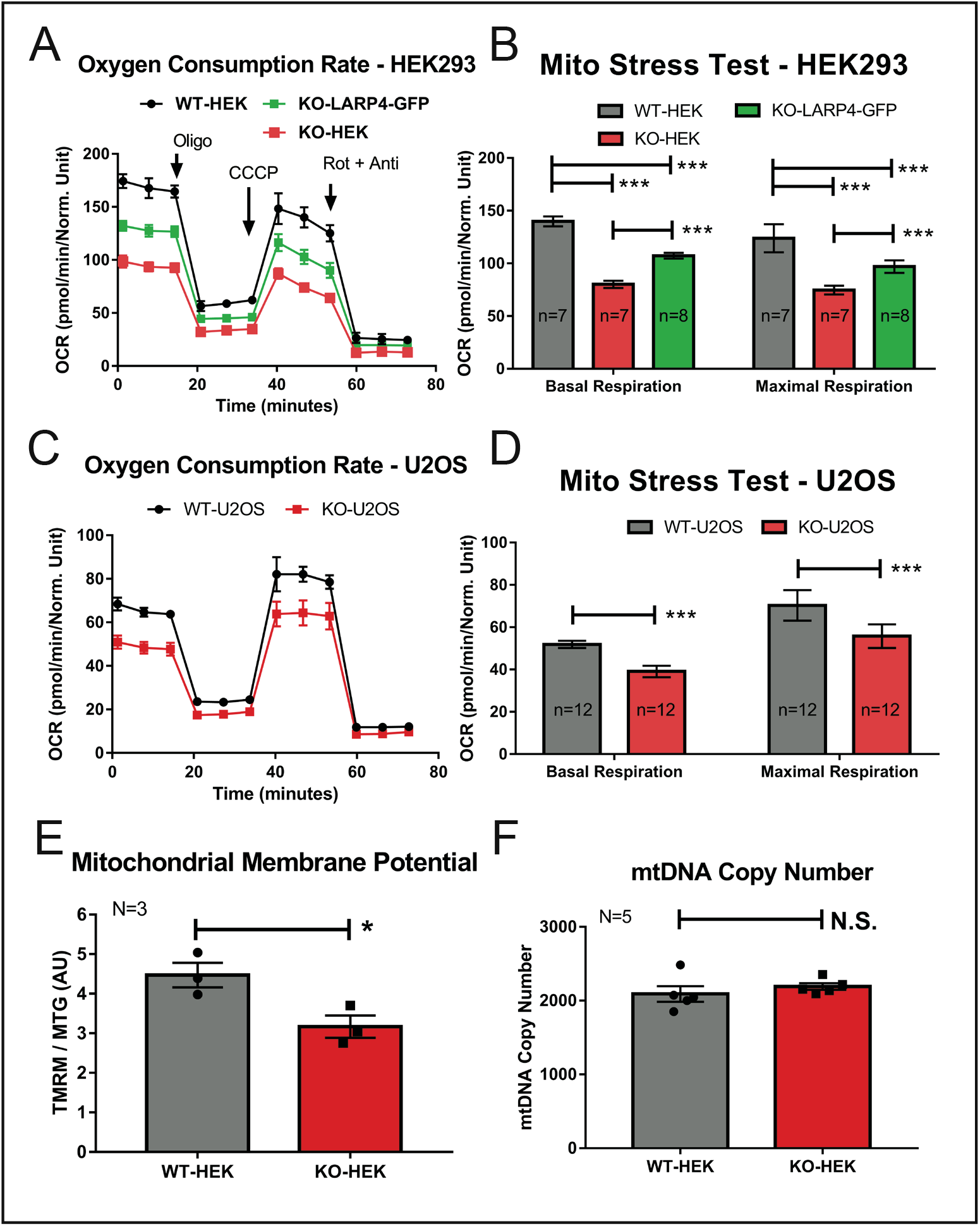
LARP4 promotes oxidative phosphorylation function. (A-B) Seahorse extracellular flux analysis of oxygen consumption rates (OCR) from wild-type HEK293 cells, HEK293^LARP4-/-^ KO cells and HEK293^LARP4-/-^ KO-LARP4-GFP Rescue cells. Basal and maximal respiration rates (B) are calculated from the changes in OCR (A) in response to inhibitor addition. (C-D) Seahorse extracellular flux analysis of oxygen consumption rates (OCR) from wild-type U2OS cells and U2OS^LARP4-/-^ KO cells. (E) Analysis of mitochondrial membrane potential by flow cytometry of wild-type HEK293 cells and HEK293^LARP4-/-^ KO cells stained with the potential dependent dye TMRM and the potential independent dye MitoTracker Green. Averages shown are from three independent experiments (N=3). (F) Analysis mitochondrial mass by proxy using qPCR measurements of mitochondrial DNA copy number from wild-type HEK293 cells and HEK293^LARP4-/-^ KO cells samples. Averages shown are from samples from five independent experiments (N=5). See also Figures S4 and S5.

### Mitochondrial Membrane Potential Disruption

The effect of reduced OXPHOS rates on the mitochondrial membrane potential (MMP) was also explored in LARP4-deficient cells. Flow cytometry analysis on the HEK293^LARP4-/-^ KO cells stained with both TMRM (MMP dependent) and Mitotracker green (Mitochondrial mass dependent, MMP independent) showed a significantly reduced MMP as measured by the ratio of TMRE staining and Mitotracker green staining (Figure 5G). Analysis of mtDNA levels was also performed as a measure of mitochondrial mass. This analysis found no significant difference in mtDNA levels between the HEK293^LARP4-/-^ KO cells and wild type cells (Figure 5F). Indicating that significant changes in mitochondrial mass in HEK293^LARP4-/-^ KO cells is unlikely.

### LARP4 Promotes Mitochondrial Associated Translation

Because some of the protein products of the RNA targets of LARP4 were more depleted in the proteomic analysis of mitochondrial extracts compared to analysis of whole cell extracts from LARP4 depleted cells (Figure 3G), we hypothesized that reduced mitochondrial associated translation results in impaired protein targeting in HEK293^LARP4-/-^ KO cells. To assess the effect of LARP4 depletion on mitochondrial associated translation the puromycin incorporation assay was performed on mitochondrial extracts. Cells were treated transiently with puromycin before preparation of mitochondrial extracts and puromycin incorporation measured by immunoblot (Figure 6B). These experiments showed that mitochondrial associated translation as measured by puromycin incorporation was significantly reduced in the HEK293^LARP4-/-^ KO cells relative to the wild type cells and re-expression of LARP4 ameliorated this phenotype (Figure 6A). Another prediction of the of this model is that impaired protein targeting will result in increased mislocalization of some LARP4 target mitochondrial proteins. To test this prediction, we used immunofluorescence staining analysis and high-resolution confocal imaging to measure the percent cytosolic signal (non-mitochondrial cell signal) of the nuclear-encoded mitochondrial OXPHOS protein COX7A2 in HEK293 LARP4 KO and WT cells. This protein is encoded by a LARP4 target mRNA and was also more depleted in the MITO-TMT (KO/WT=0.64) compared to the WCE-TMT (KO/WT=0.86). This analysis revealed a significantly greater non-mitochondrial COX7A2 signal present in HEK293^LARP4-/-^ KO cells (KO:76% vs WT:57% and KO:66% vs WT:47%) in two independent experiments (Figure 6C and 6D) indicating COX7A2 mislocalization. Together these observations support a model in which LARP4 promotes mitochondrial associated translation to enhance protein targeting of some of its NEMmRNA targets. This provides mechanistic insights into how LARP4 promotes OXPHOS function.

**Figure 6.**
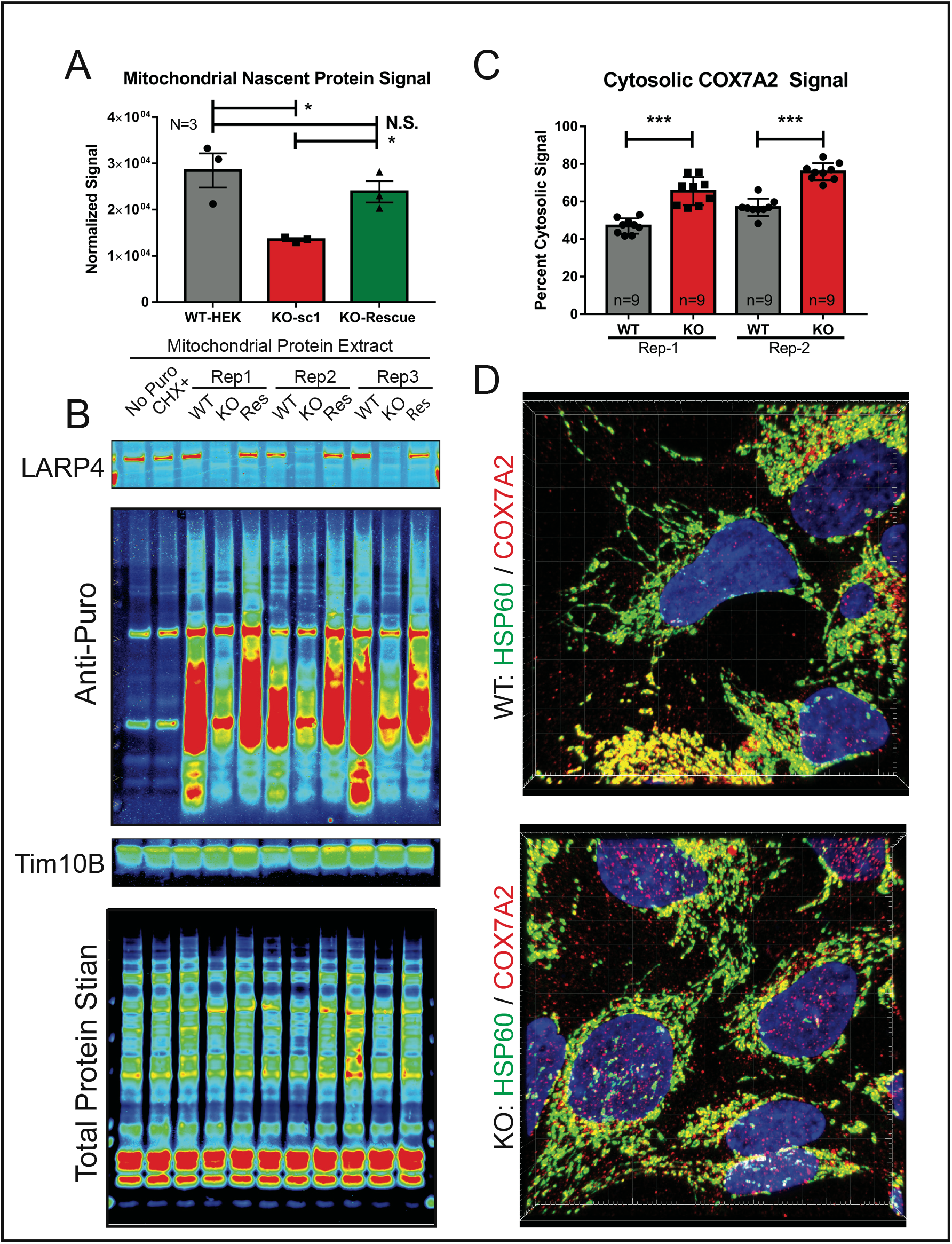
LARP4 promotes mitochondrial associated translation. (A-B) Analysis of mitochondrial associated translation rates by isolating mitochondria from puromycin treated cells and measuring puromycin incorporation. (A) Quantification of average normalized mitochondrial associated puromycin signal from biological replicates (N=3). (B) Puromycin Immunoblots used for mitochondrial puromycin incorporation assay. Controls include immunoblots for LARP4 to show depletion and TIM10B to show mitochondrial enrichment as well as a quantitative total protein stain for normalization. (C-D) Analysis of protein localization of COX7A2 (LARP4 target) by immunofluorescence staining with the mitochondrial protein HSP60 used as a mitochondrial marker and low threshold mask for cell area for image analysis. (C) Average percent cytosolic COX7A2 signal was quantified from nine fields of view (n=9) of the HEK293^LARP4-/-^ KO cells and wild-type cells from two independent biological replicates (Rep-1 and Rep-2). (D) Representative images from each condition and replicate are shown.

## Discussion

Mitochondrial gene expression presents several unique biological challenges that must be overcome to ensure mitochondrial protein homeostasis and function. The overwhelming majority of mitochondrial proteins are encoded by mRNAs transcribed from nuclear genes, all of which must be translated by cytosolic ribosomes and imported into the mitochondria by various partially redundant pathways (Bykov et al., 2020). These pathways include polypeptide dependent pathways, facilitated by the N-terminal mitochondrial targeting sequence present on many but not all mitochondrial proteins (Bolender et al., 2008), as well as mRNA dependent targeting pathways, which are facilitated by RNA-binding proteins (RBPs) that influence the abundance, spatial distribution, and translational activity of their RNA targets (Bethune et al., 2019).

Efficient targeting of proteins to the mitochondria is particularly important for certain classes of proteins. These include proteins whose ectopic expression in the cytosol has potentially deleterious effects for the cell. This has been demonstrated for both respiratory chain complex proteins (RCCPs) and mitochondrial ribosome proteins (MRPs) (Boos et al., 2019, Bykov et al., 2020, Rawat et al., 2019, Williams et al., 2014, Wrobel et al., 2014, Yoo et al., 2005). Additionally, for proteins that assemble into large multi-protein complexes, such as RCCPs and MRPs (Figure 2A), spatial and temporal coordination of translation also has the potential to promote co-translational complex assembly, a process that has been demonstrated to be beneficial in several contexts (Kamenova et al, 2019, Keil et al, 2019, Schwarz et al, 2019).

Post-transcriptional regulation of nuclear-encoded mitochondrial mRNAs (NEMmRNAs) including localization to mitochondrial membrane and localized translation is a fundamentally conserved biological process from yeast to metazoans; however, the protein factors (RBPs) and the mRNA sequences involved differ. A more complete characterization of the RBPs and their RNA interactions involved in this process in the context of human cells will facilitate greater insight into the mechanisms by which these RBPs promote efficient expression of mitochondrial proteins, potentially leading to a better understanding of the plethora of diseases where mitochondrial dysfunction is present.

In this report we elucidate a novel function of LARP4 by which it specifically binds select groups of functionally important mitochondrial mRNAs to help maintain mitochondrial protein homeostasis and facilitate proper oxidative phosphorylation function. Our results provide direct evidence that LARP4 binds mRNAs encoding RCCPs and MRPs and implicate LARP4 in positively regulating the expression of these proteins. Additionally, we demonstrate that LARP4 is required for normal oxidative phosphorylation rates.

We found that the RNA-binding protein LARP4 is required for normal translation rates, consistent with previous reports linking LARP4 to polysome stability (Yang et al., 2011). Later studies demonstrated LAPP4 to have a functional role in maintaining poly-A tail (PAT) length across the transcriptome via competition with deadenylases (Mattijssen et al., 2017, and Mattijssen et al., 2020). Interestingly, LARP4-mediated PAT length maintenance was observed across the transcriptome, except for a subset of NEMmRNAs where LARP4 did not appear to play a significant role in maintaining PAT length (Mattijssen et al., 2020). A possible explanation for this, made possible by our results, is a model in which direct RNA-targets of LARP4, such as NEMmRNAs, are bound by LARP4 in way that does not reduce deadenylase activity, whereas indirect LARP4 RNA-targets interact with LARP4 in a way that leads to reduced deadenylase activity, possibly through competing with deadenylases for protein-protein interactions with PABPs.

Another recent study described the recruitment of LARP4 to the surface of mitochondria by the protein kinase A adaptor protein AKAP1 to facilitate local translation of AKAP1 associated mRNAs (Gabrovsek et al., 2020). Our results extend beyond this study by showing that LARP4 functions not only as an RNP scaffolding factor involved in enhancing mitochondrial associated translation of AKAP1 associated mRNAs but also as a RBP with its own specific NEMmRNA targets that are translated in a LARP4 dependent manner. Additionally using a similar assay as Gabrovsek et al., 2020, we show that LARP4 depletion reduces mitochondrial associated translation, phenocopying the effect of AKAP1 depletion on mitochondrial associated translation. The mitochondrial associated puromycin translation assay we used only gives information about global mitochondrial associated translation and does not provide information about the translation of individual gene products. Further technical advances enabling the measurement of localized translation of specific gene products in mammalian cells (i.e., mammalian proximity Ribo-seq) would be useful in investigating if the translation of LARP4 mRNA targets are specifically promoted in the vicinity of the mitochondria.

In their proteomic screen for AKAP1 interacting partners, Gabrovsek et al., 2020 also found PABPC1 and PABC4, two of the poly-A binding protein paralogs, as a highly enriched AKAP1 interactors. Interestingly, in our analysis of the ENCODE eCLIP datasets we found that PABC4 also has an RNA-target set highly enriched for NEMmRNAs. Future studies should investigate what role PABC4 plays in mitochondrial gene expression and if AKAP1, LARP4 and PABC4 function cooperatively.

The data presented in this study provide strong support for a model in which LARP4 binds select mitochondrial mRNAs and positively influences the abundance of their protein products through post-transcriptional processes maintained by LARP4. Because we did not observe significant changes in mRNA abundance in the LARP4 target genes studied, it is unlikely that LARP4 functions as a selective regulator of transcript stability in the context of these mRNAs. Given that recruitment of LARP4 to the cytoplasmic surface of the mitochondria has been previously demonstrated (Gabrovsek et al., 2020) and we demonstrated that mitochondrial associated translation was reduced upon LARP4 depletion, LARP4 may play a direct role in the recruitment of a subset of its target mRNAs to the vicinity of the mitochondria to enhance their local translation and mitochondrial protein targeting. Additionally, both our proteomic analysis and immunofluorescence analysis of LARP4-depleted cells demonstrated non-mitochondrial accumulation of protein products encoded for by mitochondrial RNA targets of LARP4, implicating LARP4 in the process of mitochondrial protein targeting. Together these results are consistent with LARP4 promoting OXPHOS function by enhancing mitochondrial associated translation to facilitate efficient protein targeting to the mitochondria.

## Acknowledgements

We thank Dr. Eric Van Nostrand for initial guidance on the project. B.M.L. and this work were supported by NIH T32 Cell and Molecular Genetics Training Program (GM007240) as well as grants and awards from NIH to T.H. (CA 080100, CA 082683 and CA 242443). This work was supported by grants from the NIH to G.W.Y. (U41 HG009889 and HG004659). G.W.Y. is supported by an Allen Distinguished Investigator Award, a Paul G. Allen Frontiers Group advised grant of the Paul G. Allen Foundation. T.H. is a Frank and Else Schilling American Cancer Society Professor and holds the Renato Dulbecco Chair in Cancer Research. This work was supported by the Mass Spectrometry Core of the Salk Institute with funding from NIH-NCI CCSG: P30 014195, and the Helmsley Center for Genomic Medicine. We thank J. Diedrich and A. Pinto for proteomics technical support. We thank Dr. Sandy Mattijssen and Dr. Richard J. Maraia for providing a LARP4 antibody.

## Author contributions

Conceptualization, B.M.L., G.W.Y. and T.H.; Investigation, B.M.L.; Vector cloning, B.M.L. and C.Y.C.; Formal analysis and Visualization, B.M.L.; metagene analysis, B.M.L. and H.L.H.; Writing – Original Draft, B.M.L.; Writing – Review and Editing, B.M.L., G.W.Y. and T.H.; Funding Acquisition, G.W.Y. and T.H.

---

**Figure S1.**
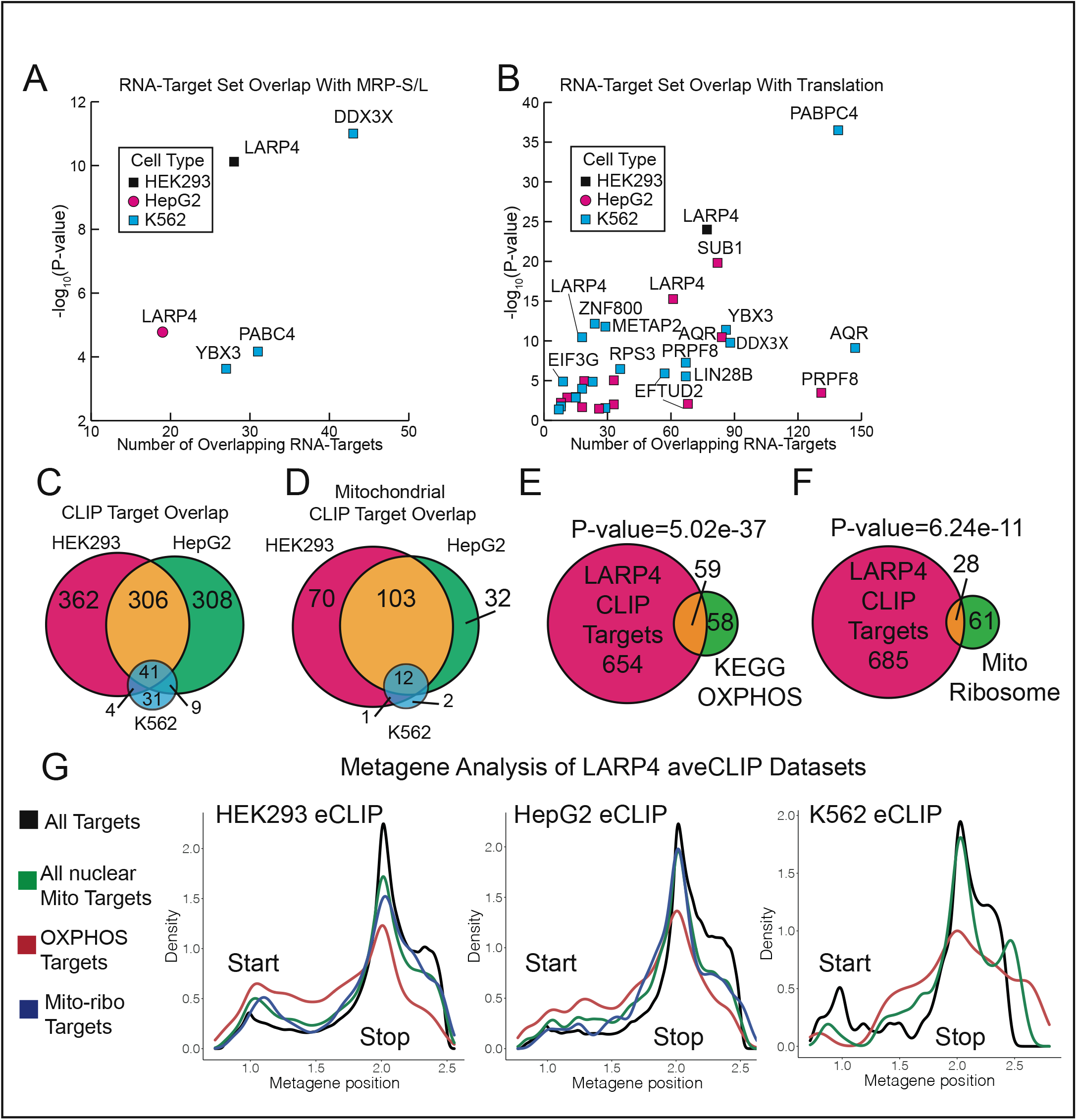
LARP4 has a preference for binding nuclear-encoded mitochondrial mRNAs. Related to Figure 1. (A) Enrichment analysis results for RNA-target set (eCLIP data) overlap with mitochondrial ribosome proteins (MRP-S/L) defined by gene ontology term GO_0005761. Only RBPs with significant overlap (p-value<0.05) are shown. (B) Enrichment analysis results for RNA-target set (eCLIP data) overlap with proteins in the Translation pathway (R_HSA_72766). Only RBPs with significant overlap (p-value<0.05) are shown. (C) Venn diagram showing the overlap between gene-target list of each of the three LARP4 eCLIP datasets from HEK293, HepG2, K562 cells. (D) Venn diagram showing the overlap between gene-targets that are also mitochondrial genes. (E) Venn diagrams showing the overlap HEK293’s LARP4 eCLIP gene-target list with oxidative phosphorylation genes (HSA_00190) (F) Venn diagrams showing the overlap HEK293’s LARP4 eCLIP gene-target list with mitochondrial ribosome proteins (GO:OOO5761) (G) Metagene analysis of eCLIP-seq data from HEK293 cells, HepG2 cells and K562 cells. Peak density is plotted across all genes (black), nuclear-encoded mitochondrial genes (green), OXPHOS genes (red) and mitochondrial ribosome genes (blue). Gene-target lists of each CLIP-seq data set are defined as genes encoding an mRNA containing at least one CLIP peak that passes significance thresholds (-log10Pvalue>7, log2foldchange (IP/input)>4).

**Figure S2.**
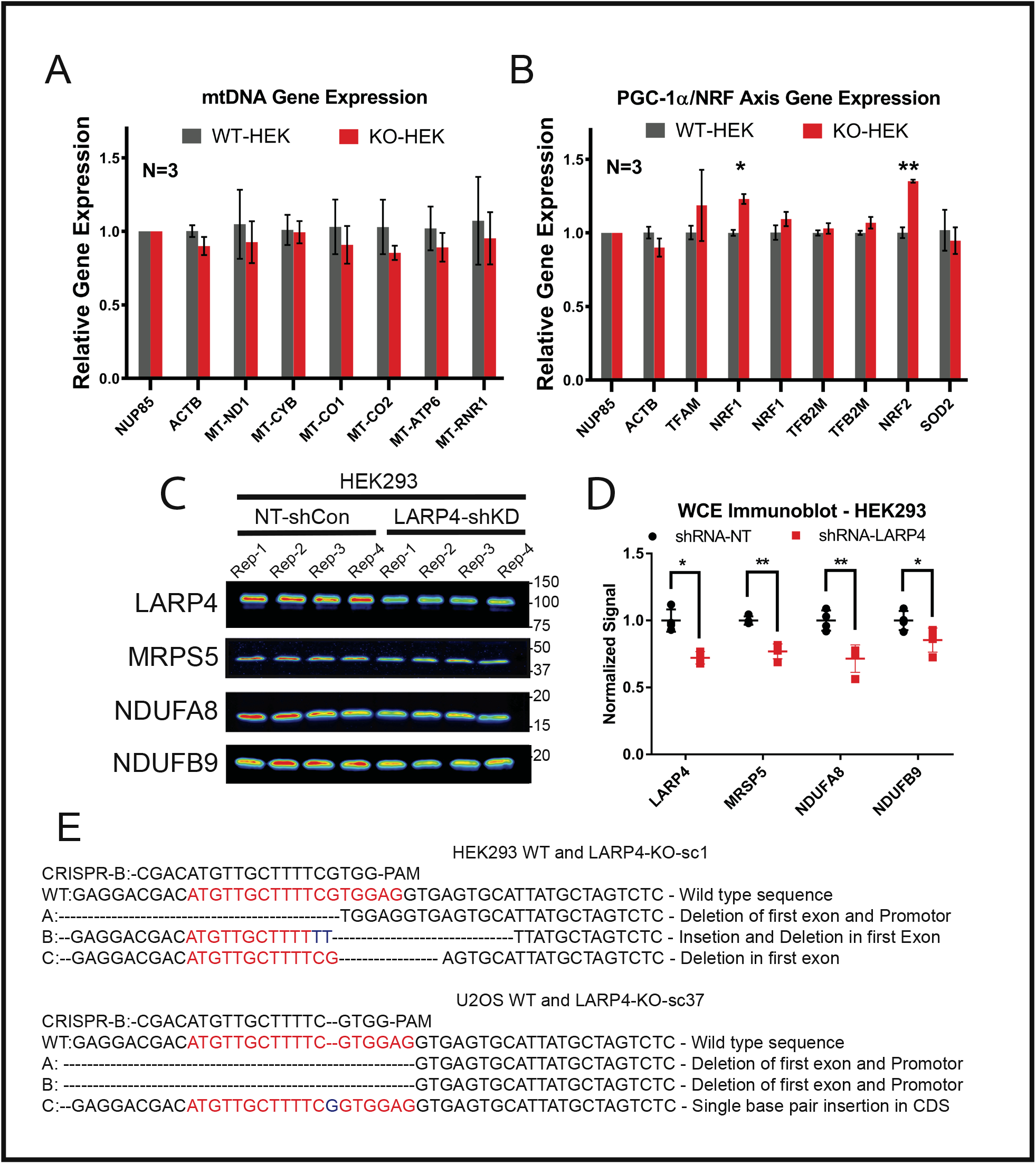
Analysis of mRNA abundance and CRISPR cut site analysis. Related to Figure 2. (A-B) Analysis of mRNA abundance in the HEK293 KO/WT cells by qPCR using probes targeting genes expressed from the mitochondrial DNA (A) or genes involved in the PGC-1alpha/NRF pathway (B). (C) Immunoblot analysis of HEK293 cells expressing either an shRNA targeting LARP4 (LARP4-shKD) or a non-targeting shRNA (NT-shCon). (D) Quantification of biological replicates shown in immunoblot panels. Band intensities are normalized by a quantitative total protein stain. (E) Analysis of the LARP4 genomic region flanking the gRNA target sites by Sanger sequencing. Unique sequences recovered for each line are shown aligned to the wild-type sequence. Insertions are shown in blue, deletions are indicated with a dash and the CRISPR target site in red.

**Figure S3.**
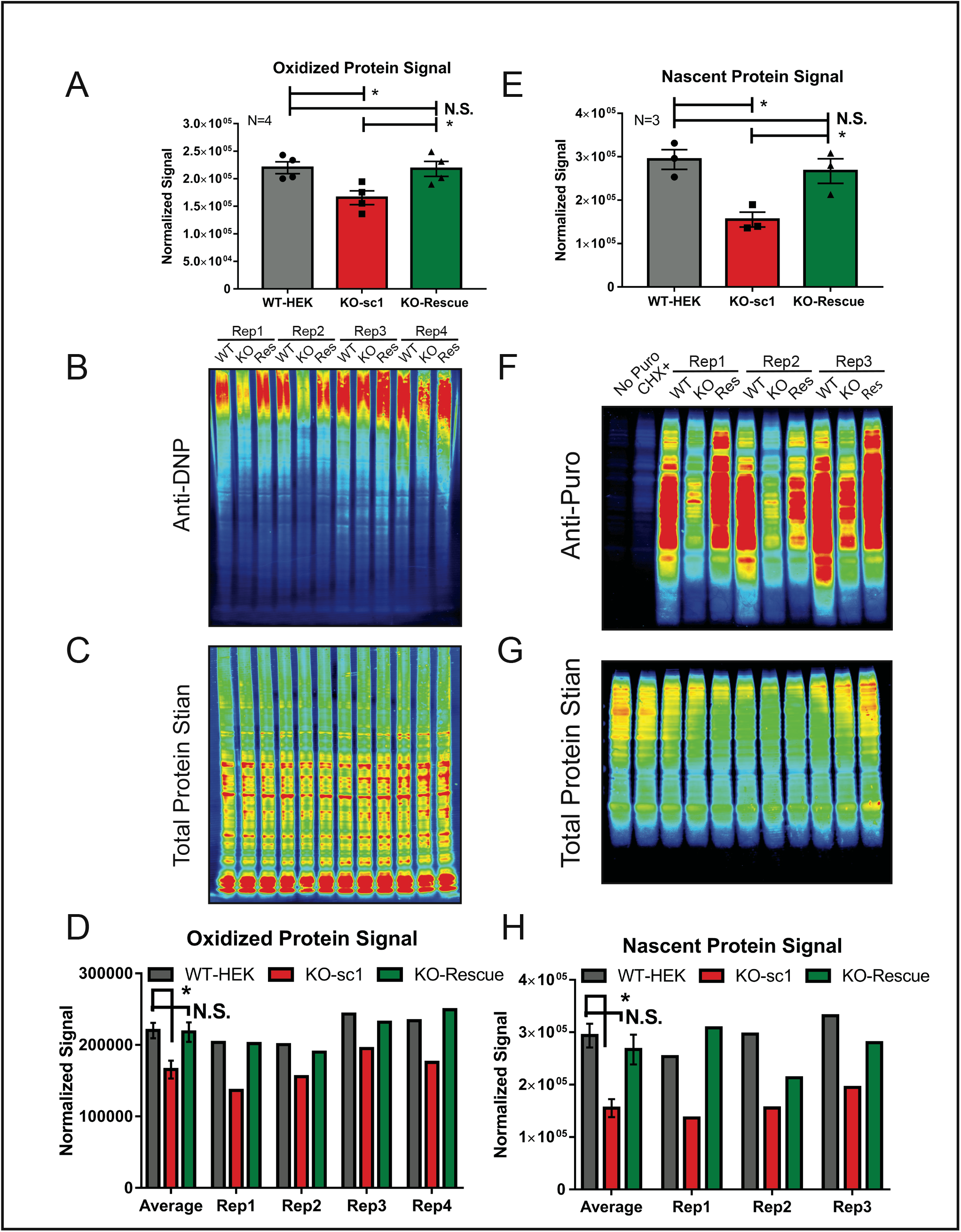
Oxiblot assay and puromycin assay. Related to Figure 4. (A) A summary plot of average (N=4) normalized oxidized protein signal from biological replicates. (B) An anti-DNP immunoblot probing for the derivatized oxidized proteins. (C) A quantitative total protein stain of the blot in panel B. (D) A bar graph showing the normalized oxidized protein signal for each of the replicates. (E) A summary plot of average (N=3) normalized puromycin incorporation signal from biological replicates. (F) An anti-Puromycin immunoblot probing for the nascent proteins terminated by puromycin. (G) A quantitative total protein stain of the blot in panel B. (H) A bar graph showing the normalized puromycin incorporation signal for each of the replicates.

**Figure S4.**
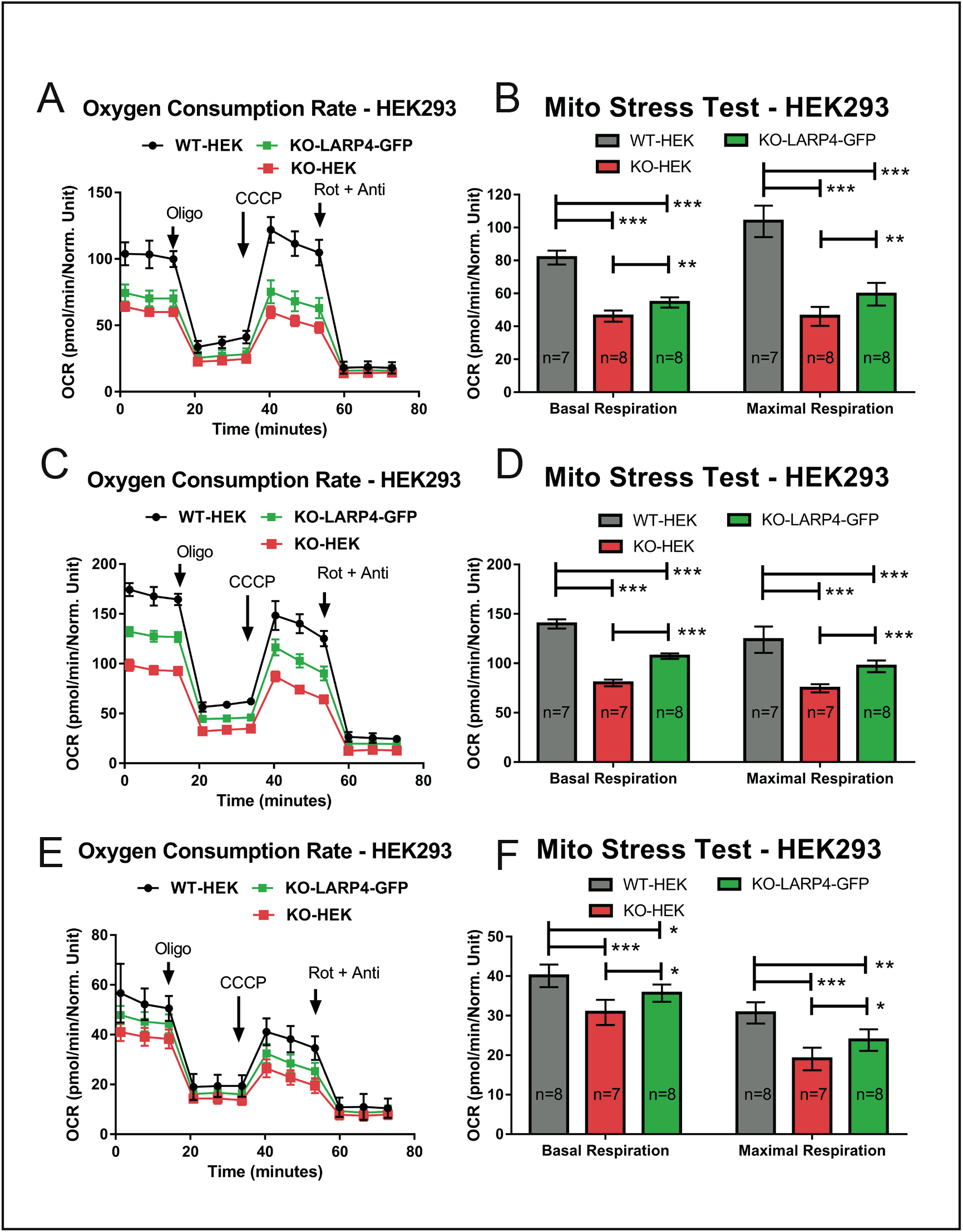
Replicate data for Seahorse Mito Stress Test on HEK293 LARP4 KO cell line. Related to Figure 5. (A-B) Replicate 1. (C-D) Replicate 2. (E-F) Replicate 3.

**Figure S5.**
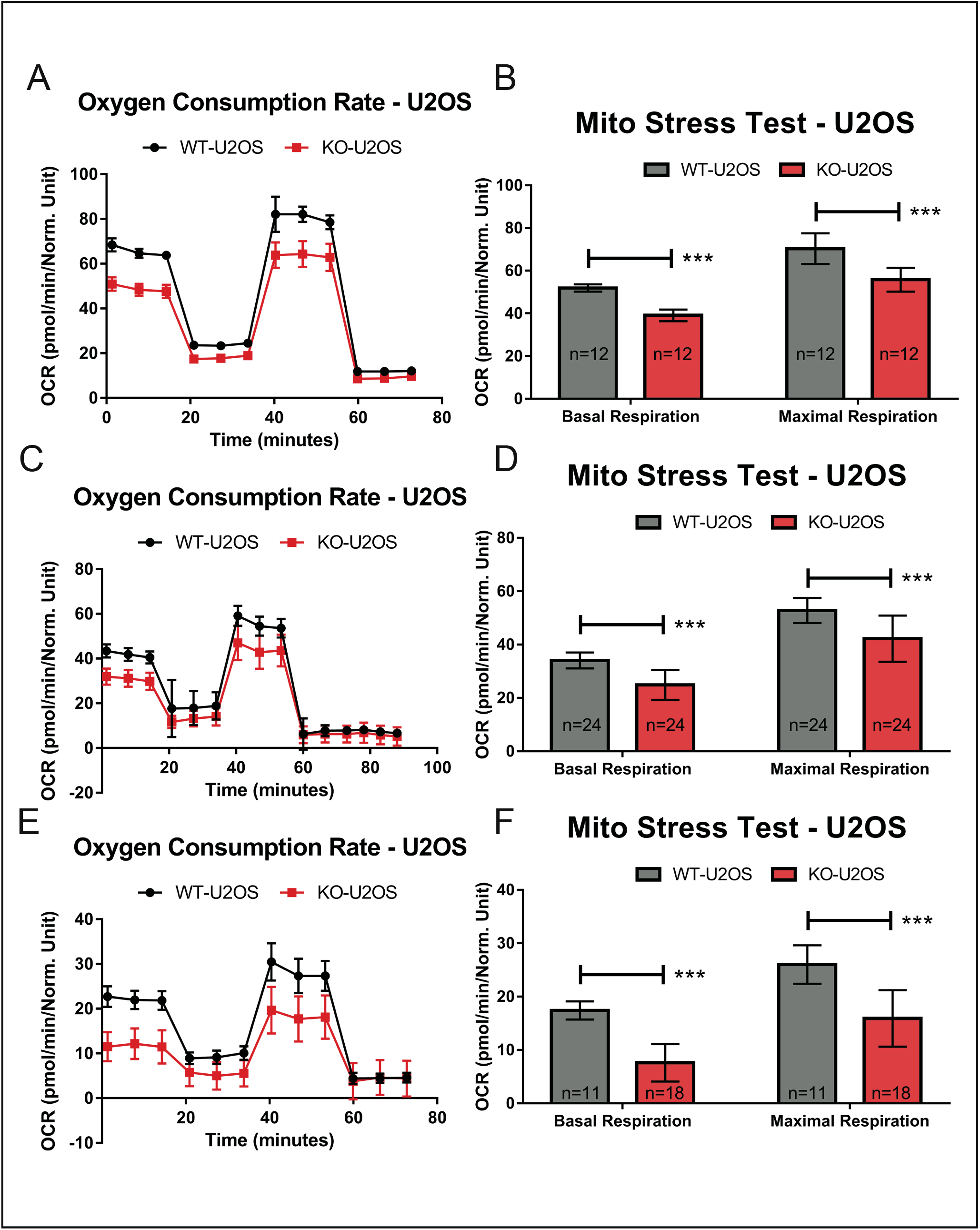
Replicate data for Seahorse Mito Stress Test on U2OS LARP4 KO cell line. Related to Figure 5. (A-B) Replicate 1. (C-D) Replicate 2. (E-F) Replicate 3.

## Materials and Methods

### DNA Constructs

The coding region of the full length LARP4 was amplified from pFLAG-CMV2 plasmid containing a LARP4 expression cassette (gift from the Richard Maraia lab) and cloned into a gateway entry plasmid by Gibson assembly. Subsequently lenti-viral expression plasmids were produced via gateway cloning. CRISPR plasmids were cloned according to Zhang lab protocols (Ann Ran et al. 2013).

### Cell culture and manipulations

All cell lines were maintained in DMEM (Corning,10-013-CV) supplemented with 10% fetal bovine serum at 37 °C with 10% CO_2_. Cultures were passaged with accutase (Innovative Cell Technologies, AT104) every three days. HEK293 and U2OS cell lines were obtained from ATCC and cultured for 30 passages or less. Periodic testing for mycoplasma was performed. All transfections were performed using lipofectamine 2000 (Thermo-Fisher,11668030) according to the manufacturer’s instructions. All transductions were performed using filtered cell culture media from cells transfected with lenti-viral production plasmids.

### CRISPR knockout cell line generation

LARP4 knock cell lines were generated with CRISPR/Cas9 using a two-guide transfection strategy. Cas9 and sgRNA guides were introduced into cells by transfection of plasmids expressing Cas9 and sgRNA guides. Forty-eight hours after transfection, cells were singularized and sorted into individual wells by flow cytometry to generate subclone cultures. Recovered clones were first screened by PCR using primers flanking the two adjacent CRISPR target site. Clones showing evidence of a deletion in the target region were selected for validation by immunoblot analysis. CRISPR guides sequences and PCR screening primers were used and are contained in the key recourses table. For the KO clones used in this study, the genomic region flanking the gRNA target site was amplified and cloned into a cloning vector and plasmids from 10 individual transformants for each clone sent for sequencing.

### Incucyte cell proliferation assays

For analysis of cell proliferation rates, the Zoom Incucyte live cell imaging device was used to make image-based cell density measurements of cell cultures at regular intervals. The manufacturer’s image analysis software was used to calculate either percent well confluency (HEK293) or direct cell counts (U2OS) at each time point. These cell density measurements were used to calculate density doubling times for individual wells of the cell culture plate. For experiments with HEK293 cells 6,000 cells were plated into wells of 96 wells plates, 24 hours prior to transfer into the Incucyte device and growth measurements recorded for 4-5 days. Reported confluency doubling times represent averages of three biological replicates, each of which represents an average of technical replicates, which consisted of eight individual wells. For experiments with U2OS cells, cells were first transduced with a lentivirus expressing a nuclear red fluorescent protein to directly track cell counts. For experiments with U2OS cells only two biological replicates were performed and cell doubling times reported are averages of technical replicates.

### Immunoblotting

For standard immunoblot analysis, lysates were prepared in RIPA lysis buffer (Thermo-Fisher,89900) supplemented with EDTA-free complete protease inhibitors (Sigma,5892791001). Protein concentrations determined by the BCA assay (Thermo-Fisher,23227). Samples concentrations were normalized and prepared for PAGE in a reducing loading buffer, depending on the target protein 10-30 ug of protein per sample were resolved through acrylamide gels. Proteins were then transferred onto PVDF membranes (Millipore, IPFL00010) via the wet-transfer procedure. Total protein transferred protein was then quantified using the Revert fluorescent total protein stain (Li-Cor,926-11011) according to the manufacturer’s instructions. Membranes were then blocked via a one hour incubation in blocking buffer (150mM NaCl, 50mM Tris, pH 7.5, 5% w/v bovine serum album). Primary antibody incubations were carried out for two hours at room temperature in blocking buffer supplemented with 0.1% Tween-20. Secondary antibody incubations were carried out for one hour at room temperature in blocking buffer supplemented with 0.1% Tween-20 and 0.01% SDS. The following primary antibodies were used at 1:1000 dilution unless stated otherwise: (LARP4, Maraia Lab), (NDUFA8, Abcam, ab184952, dilution: 1:20,000), (NDUFB9, Abcam, ab200198, dilution: 1:5,000), (COX5B, Abcam, ab180136, dilution: 1:10,000), (COX6B1, Abcam, ab131277), (MRPS5, Abcam, ab96291), (MRPL24, Santa Cruz Biotechnology, sc-393857, dilution: 1:100), (NDUFA1, Abcam, ab176563), (ATP5L, Invitrogen, PA5-60783), (Calnexin, Santa Cruz Biotechnology, sc-11397, dilution: 1:100), (TBP, Cell Signaling, 44059), (TIMM10B, Invitrogen, PA597081), (COX7A2, Proteintech, 18122-1-AP, dilution 1:100), (Anti-puromycin, Millipore, MABE343, dilution: 1:20,000), (Anti-DNP, Millipore, S7150, dilution: 1:150)(need to add COX7A2 antibodies used for IF). The following secondary antibodies were used at a 1:20,000 dilution: (Goat anti-rabbit 800 nm, Invitrogen, SA535571), (Goat anti-mouse 800 nm, Invitrogen, SA535521), (Goat anti-rabbit Alexa flour 680, Invitrogen, A-21109), (Goat anti-mouse Alexa flour 680, Invitrogen, A-21058). After each antibody incubation membranes were washed three times in wash buffer (150 mM NaCl, 50 mM Tris, pH 7.5, 0.1% Tween-20) for five min. Stained membranes were rinsed in TBS (150 mM NaCl, 50 mM Tris, pH 7.5) and visualized using the Odyssey imaging system. The LI-COR imaging software was used to quantify both total transferred protein per lane and signal intensity of the target band. All reported immunoblot signal quantifications are the target band signal normalized by the total protein signal of the lane containing the target band.

### Oxiblot assay

For analysis of oxidized protein abundance using the Oxiblot assay kit (Millipore, S7150), lysates were prepared in RIPA lysis buffer supplemented with 50 μM DTT. Protein concentrations between samples were normalized and oxidized proteins were then derivatized with DNP according to the manufacturer’s instructions. Immunoblotting was then performed and quantified as previously described, using the anti-DNP primary antibody provided with Oxiblot assay kit (Millipore, S7150).

### Puromycin incorporation assay

For analysis of translation rates by the puromycin incorporation assay (Schmidt, et al. 2009), cells in culture were incubated in the presence of puromycin (10 μg/ml) for 10 min prior to lysate collection in RIPA buffer. Immunoblotting was then performed and quantified as previously described using an anti-puromycin primary antibody (Millipore, MABE343). Negative control sample cell cultures were processed similarly with the exception that cells were either incubated with unmodified media, or puromycin labeling media supplemented with cycloheximide at a final concentration of 100 μg/ml.

### RT-qPCR analysis

For analysis of mRNA abundance, total RNA extracts were collected from cell cultures containing approximately 500,000 cells by directly applying Trizol reagent (Thermo-Fisher, 15596026). Total RNA was further purified using the Direct-zol column purification kit (Zymo, R2050). Reverse transcriptase reactions were carried out using the SuperScript™ III Reverse Transcriptase kit (Invitrogen, 18080044), according to the manufacturer’s instructions. A qPCR analysis was then performed on the resulting cDNA with Power SYBR™ Green PCR Master Mix (Thermo-Fisher, 4368577) using primer pairs listed in the key resource table. For each sample type three biological replicates (independent collection days) were assayed, with at least two technical replicates (replicate qPCR reactions) performed for each biological replicate.

### mtDNA abundance analysis

For analysis mtDNA abundance, approximately 500,000 cells were collected and pelleted. Cell pellets were then resuspended in 50 mM NaOH and incubated at 95 °C for one hour. Samples were then neutralized with 1 M Tris, pH 8.0 using 1/10^th^ the volume of lysis buffer used. DNA concentration was then estimated by spectrophotometry and qPCR performed using 3 ng of DNA per 10 μl reaction of Power SYBR™ Green PCR Master Mix (Thermo-Fisher, 4368577). For each sample two reactions were prepared using primers targeting either the nuclear DNA (beta-2-microglobulin) or primers targeting the mitochondrial DNA (mitochondrially encoded tRNA leucine 1). The ratio of cT values from the two primer sets was used to estimate the ratio of mitochondrial DNA to nuclear DNA. For each sample type at least three biological replicates (independent collection days) were assayed, with at least two technical replicates (replicate qPCR reactions) performed for each biological replicate.

### Magnetic Isolation of mitochondria

Magnetic isolation of mitochondria was performed using the Militenyi Biotec Human Mitochondria Isolation Kit according to the manufacturer’s instructions with additional modifications and the following experiment specific details. Hek293 cells were seeded at a density of 15X10^6 per 15 cm plate, 24 hours before collection. Cells were harvested using DPBS and a rubber policeman, approximately 10% of the cells collected were pelleted and set aside to be used as matched whole cell extract samples and the remaining 90% pelleted and resuspend in 1ml of the kit’s lysis buffer, which was supplemented with protease inhibitors (Sigma, 5892791001). A 26-gauge needle and 1ml syringe were used to homogenize the cell lysates. After optimization, the following protocol was found to be effective for 85-95% cell disruption: 5 strokes of the entire homogenate volume, ice for 60 seconds then repeat three more times for a total of 20 stroke repetitions. The final pellet for the mitochondria extracts and the whole cell extracts were resuspended in RIPA buffer supplemented with protease inhibitors (Sigma, 5892791001). To remove the antibody-conjugated magnetic nanobeads prior analysis by quantitative TMT mass spectrometry, lysed mitochondria extracts were passed over fresh LS columns from the isolation kit, that had been equilibrated with RIPA buffer.

### TMT-mass spectrometry and analysis

Samples were precipitated by methanol/ chloroform and redissolved in 8 M urea/100 mM TEAB, pH 8.5. Proteins were reduced with 5 mM tris(2-carboxyethyl)phosphine hydrochloride (TCEP, Sigma-Aldrich) and alkylated with 10 mM chloroacetamide (Sigma-Aldrich). Proteins were digested overnight at 37 ^o^C in 2 M urea/100 mM TEAB, pH 8.5, with trypsin (Promega). The digested peptides were labeled with 10-plex TMT (Thermo, 90309), pooled samples were fractionated by basic reversed phase (Thermo, 84868).

The TMT labeled lysate samples were analyzed on a Fusion Lumos mass spectrometer (Thermo). Samples were injected directly onto a 25 cm, 100 μm ID column packed with BEH 1.7 μm C18 resin (Waters). Samples were separated at a flow rate of 300 nL/min on an EasynLC 1200 (Thermo). Buffer A and B were 0.1% formic acid in water and 90% acetonitrile, respectively. A gradient of 1–25% B over 180 min, an increase to 40% B over 30 min, an increase to 100% B over another 20 min and held at 90% B for a 10 min was used for a 240 min total run time.

Peptides were eluted directly from the tip of the column and nanosprayed directly into the mass spectrometer by application of 2.8 kV voltage at the back of the column. The Lumos was operated in a data dependent mode. Full MS1 scans were collected in the Orbitrap at 120k resolution. The cycle time was set to 3 s, and within this 3 s the most abundant ions per scan were selected for CID MS/MS in the ion trap. MS3 analysis with multinotch isolation (SPS3) was utilized for detection of TMT reporter ions at 60k resolution (Graeme C. McAlister et al., 2014). Monoisotopic precursor selection was enabled and dynamic exclusion was used with exclusion duration of 10 s.

Protein and peptide identification were done with Integrated Proteomics Pipeline – IP2 (Integrated Proteomics Applications). Tandem mass spectra were extracted from raw files using RawConverter (Lin He et al., 2015) and searched with ProLuCID (T. Xu et al., 2015) against Uniprot human database. The search space included all fully-tryptic and half-tryptic peptide candidates. Carbamidomethylation on cysteine and TMT on lysine and peptide N-term were considered as static modifications. Data was searched with 50 ppm precursor ion tolerance and 600 ppm fragment ion tolerance. Identified proteins were filtered to using DTASelect (David L. Tabb et al., 2002) and utilizing a target-decoy database search strategy to control the false discovery rate to 1% at the protein level (Peng et al., 2003). Quantitative analysis of TMT was done with Census (Sung Kyu Robin Park et al., 2014) filtering reporter ions with 20 ppm mass tolerance and 0.6 isobaric purity filter.

Protein total intensity on each channel was normalized by the sum of all proteins in the same channel. These normalized intensity values for each replicate (N=4) were averaged together by group (WT or KO) and used to calculate average normalized protein abundance for each protein. These values were used to calculate KO/WT ratios for each protein and determine sets of proteins present in increased or decreased abundance for gene ontology analysis. Common keratin contaminates were removed manually.

### Extracellular flux analysis

Oxygen consumption rates and proton efflux rates were measured using the XF-96 extracellular flux analyzer (Agilent) and XF-96 FluxPaks (Agilent). The mito-stress test (MTS) assay was carried out according to the manufacturer’s instructions with additional modifications and the following experiment specific details. Approximately 24 hours prior to assay, cells were seeded onto XF-96 cell culture plates and at a density of 25,000 or 16,000 cells per well for HEK293 cells or U2OS cells respectively. For experiments with HEK293 cells the following final concentrations of small molecules were applied: oligomycin (1.5 μM), CCCP (1.0 μM) Rotenone (0.5 μM) and Antimycin A (0.5 μM). For experiments with U2OS cells the following final concentrations of small molecules were applied: oligomycin (1.0 μM), CCCP (2.0 μM), Rotenone (0.5 μM) and Antimycin A (0.5 μM). The following small molecules required for the MTS assay were ordered individually and 10 mM stocks prepared and stored at −20°C: (CCCP, Sigma, C2920), (Oligomycin, Sigma, O4876), (Antimycin A, Sigma, A8674). The assay media used was XF DMEM base media (Agilent, 103575-100) supplemented with glucose 10 mM, pyruvate 1 mM and glutamine 2 mM. For MTS assay well normalization, DAPI (Thermo-Fisher, D3571) was added to the final small molecule injection solution such that final well concentration was 1 μg/ml. After completion of flux measurements, the cell culture plates were transferred to the Celigo plate cytometer (Cyntellect), DAPI stained nuclei imaged, and direct cell counts for each well made using the Celigo cytometer software. For analysis of extracellular flux measurements, the Wave Desktop program (Agilent) was used.

### Flow Cytometry

For flow cytometry measurements, cells were seeded 24 hours prior to collection at a density of 250,000 cells per well of a 12-well plate. Cells were harvested with accutase, pelleted, and resuspend in flow buffer (DPBS, 2% FBS, 1 μg/ml DNase I, 1 μg/ml DAPI) prior to flow analysis. Cell suspensions were analyzed with a LSR II flow cytometer (BD Biosciences) and data analyzed with FACSDiva software (BD Biosciences). For analysis of mitochondrial membrane potential (MMP), cells were co-stained with the MMP dependent dye, tetramethylrhodamine-methyl-ester-perchlorate (TMRM) (Thermo-Fisher, T668) and the mitochondrial mass dependent dye, MitoTracker™ Green (MTG) (Thermo-Fisher, M7514). MMP measurements were then normalized by the mitochondrial mass dependent dye measurements. For flow cytometry staining, media was replaced with staining media containing TMRM (50 nM) and MTG (50 nM) and cultures incubated at 37 °C for 25 min prior to harvest. To validate membrane potential of TMRM dye, control samples were created by adding either a depolarizing agent (CCCP 30 μM) or a hyperpolarizing agent (oligomycin 10 μM) to the staining media.

### seCLIP-library preparation and analysis

For analysis of RNA-targets bound by LARP4 in HEK293 cells, seCLIP experiments were carried as previously described (Van Nostrand et al., 2017) with the following experiment specific details. UV-crosslinked LARP4 was with immunoprecipitated with anti-LARP4 rabbit polyclonal antibody (Bethly, A303-900A). Libraries were sequenced as 75 base-pair single-end reads on the HiSeq4000 to approximately 20 million reads. Reads were processed through the previously described eCLIP0v0.40 pipeline (Van Nostrand et al., 2017). Input normalized peaks were further filtered by the irreproducible discovery rate (IDR) and merged using merge_peaks-v0.05 ([IDR] https://github.com/YeoLab/merge_peaks).

To find specific pathway genes, we combined annotations from the MitoCarta database with Gencode annotations v19 to filter IDR peaks in specific gene groups. Nuclear encoded mitochondrial genes are defined as all genes in MitoCarta database that is not encoded on chrM. Mitochondrial encoded ribosomes are defined as any gene name within MitoCarta that contains RPS/RPL. OXPHOS is defined with the pathway annotation in MitoCarta. The filtered IDR peaks are subsequently fed into the MetaPlotR pipeline and generated the metagene plots. For motif analysis, we define a set of strong binding sites with (-log10Pvalue > 7, log2foldchange (IP/input) > 4. Along with the gene groups defined above, we ran HOMER motif analysis using our in house-pipeline (https://github.com/YeoLab/clip_analysis_legacy) for each set of IDR peaks. Briefly, peaks were stratified by regions (CDS, UTR, introns), then HOMER was used de novo to find motif with a GC-matched, region matched background sequences. We found no significant motif in any of the region or gene groups.

### Gene ontology analysis

For all gene ontology analysis, the Metascape gene ontology method and sever was used (Zho u et al., 2019). From each dataset two gene sets were generated, a set of foreground genes and a background gene set. For analysis of LARP4 eCLIP datasets (HEK293, HepG2 and K562), the set of foreground genes (genes that encode RNA-targets of LARP4) were generated from the annotated bed file of the merged IDR peaks. Genes with a CLIP peak passing the following filters were included in each set of foreground genes: log2 fold change (CLIP-IP/Input) greater than 4 and -log10(pValue) greater than 7. For background gene sets, the matched CLIP input datasets were used, all genes with at least 5 reads in any exonic region were included in each background set. For analysis of the TMT mass spectrometry datasets foreground gene sets were generated from proteins present in either increased abundance or decreased abundance and background gene sets defined by all proteins detected in each mass spectrometry dataset (Mitochondrial extracts or Whole cell extracts). Proteins with significantly increased abundance were defined as having a log2 fold change (KO/WT) of greater than 0.2 and pValue of less 0.05. Proteins with significantly decreased abundance were defined as having a log2 fold change (KO/WT) of less than negative 0.2 and pValue of less 0.05.

### Statistical analysis

Average values were tested for statistical differences using two-tailed unpaired Student’s t-tests. Averages of biological replicates were plotted as mean values ± standard error and replicate number denoted with a capital N. Averages of technical replicates were plotted as mean values ± standard deviation and replicate number denoted with a lowercase n. The statistical significance of differences of averages are indicated by * p≤0.05, ** p≤0.001, *** p≤0.0001, **** p≤0.00001. Statistical significance of enrichments was determined by the hypergeometric test.

